# Inferring physiological energetics of loggerhead turtle (*Caretta caretta*) from existing data using a general metabolic theory

**DOI:** 10.1101/070987

**Authors:** Nina Marn, S.A.L.M. Kooijman, Marko Jusup, Tarzan Legović, Tin Klanjšček

**Affiliations:** Rudjer Bošković Institute; Bijenička cesta 54, HR-10002 Zagreb, Croatiab; Vrije Universiteit Amsterdam De Boelelaan 1105, 1081 HV Amsterdam, Netherlands; Center of Mathematics for Social Creativity, Hokkaido University 12-7 Kita Ward, Sapporo 060-0812, Japan

**Keywords:** life cycle model, DEB theory, loggerhead turtle, Dynamic Energy Budget

## Abstract

Loggerhead turtle is an endangered sea turtle species with a migratory lifestyle and worldwide distribution, experiencing markedly different habitats throughout its lifetime. Environmental conditions, especially food availability and temperature, constrain the acquisition and the use of available energy, thus affecting physiological processes such as growth, maturation, and reproduction. These physiological processes at the population level determine survival, fecundity, and ultimately the population growth rate—a key indicator of the success of conservation efforts. As a first step towards the comprehensive understanding of how environment shapes the physiology and the life cycle of a loggerhead turtle, we constructed a full life cycle model based on the principles of energy acquisition and utilization embedded in the Dynamic Energy Budget (DEB) theory. We adapted the standard DEB model using data from published and unpublished sources to obtain parameter estimates and model predictions that could be compared with data. The outcome was a successful mathematical description of ontogeny and life history traits of the loggerhead turtle. Some deviations between the model and the data existed (such as an earlier age at sexual maturity and faster growth of the post-hatchlings), yet probable causes for these deviations were found informative and discussed in great detail. Physiological traits such as the capacity to withstand starvation, trade-offs between reproduction and growth, and changes in the energy budget throughout the ontogeny were inferred from the model. The results offer new insights into physiology and ecology of loggerhead turtle with the potential to lead to novel approaches in conservation of this endangered species.

## I. Introduction

Seven known species of sea turtles currently inhabit the world’s oceans. All seven are listed in the IUCN list of endangered species [1] and face various threats despite conservation measures [2]. The conservation of sea turtles is complicated by a lack of understanding of their physiology and ecology, and by a long and complex life cycle, spanning multiple habitats over a wide geographical range [3]. Metabolic processes such as growth, maturation, and reproduction are key physiological and ecological determinants, the understanding of which is also crucial for conservation efforts. These processes are influenced by genetics [4], but also by environmental conditions, such as food availability and temperature [5, 6], that constrain the acquisition and use of energy. A way to better understand the physiology and ecology of a species is to reconstruct its energy budget using the principles of a general metabolic theory (e.g. [7, 8, 9]). Indeed, the need for an energy budget approach in the research of sea turtles was identified almost a decade ago [10].

Focusing on the loggerhead turtle and one of its largest nesting aggregations, the North Atlantic population [11], we aim to reconstruct the energy budget of this species from existing data. We begin with a brief overview of loggerhead turtle physiology and ecology. Next we explain the methodology used to develop the full life cycle model, and list the data sets used in parameter estimation. By estimating the parameter values, we establish a mapping between existing data and the loggerhead turtle energy budget. We analyze the validity of the mapping, and discuss physiological and ecological implications of the reconstructed energy budget.

### 1.1 The loggerhead turtle

Three aspects of the loggerhead turtle’s physiology and ecology impede conservation efforts. These three impeding aspects are (i) a geographically wide species distribution, (ii) long and complex ontogenetic development, and (iii) late and variable reproductive output.

Loggerhead sea turtle is a migratory species with global distribution throughout the temperate zone [1]. Individuals of this species occupy habitats ranging from cold and nutrient-sparse oceanic zones to warm and food-rich neritic zones, where some of the habitat variability is related to an ontogenetic shift [12, 13] with important implications for the energy budget. Furthermore, the wide distribution of loggerhead turtles means that populations such as the North Atlantic one span multiple jurisdictions and legislative systems with different conservation targets, methods, and ultimately success[3].

The ontogenetic development of loggerhead turtles exhibits numerous fascinating characteristics. The sex of embryos is determined by nest temperature in the second third of the embryonic development [14, 15]. Throughout its ontogeny, from hatching to ultimate size, an average loggerhead turtle increases almost 25-fold in length, and 6500-fold in body mass. Straight carapace length at hatching is 4-5 cm, while body mass is around 20 g [14]. By contrast, adults range between 90-130 cm straight carapace length and between 100-130 kg body mass [14, 16].

The average female needs 10-30 years to reach puberty [17, 18]. Reproducing every 2-3 years, females lay 4-5 clutches of over a hundred eggs each [19, 20]. The reproduction rate was found to correlate with the average sea surface temperature [21, 22], as well as the large scale environmental oscillations [23].

## 2. Methods

### 2.1 Full life cycle model of the loggerhead turtle

We use the Dynamic Energy Budget (DEB) theory [24, 25, 26] to model the full life cycle of loggerhead turtles. By relying on DEB theory, we ensure that our model is thermodynamically consistent, meaning that the conservation laws of mass and energy are strictly observed. Modeled loggerhead turtles also obey several homeostasis rules as a way of coping with sudden, unfavorable changes in the environment, especially in food availability. Metabolic rates (e.g., food assimilation, somatic maintenance, etc.) follow from scaling assumptions (concise statements of these assumptions are found below) appended with the kappa rule for allocation to soma [24, 27]. The essence of the kappa rule is that energy is divided at a fixed fraction between soma and the reproductive cells. DEB model furthermore accounts for embryonic development, where turtle eggs start as blobs of energy received from mothers. This initial energy reserve is used by the embryo to start building structure and to mature enough in order to begin feeding on an outside energy source. The basic model prescribes the rate at which mothers commit energy to reproduction. We make a step forward and convert this energy into the number of eggs as if they were produced in a continuous manner. Modeling the timing and the duration of reproductive seasons is also possible by means of species- or population-specific rules for handling the storage of energy between reproductive seasons and the conversion of stored energy into eggs during one such season.

Free ranging animals owe their mobility in large part to a better homeostatic regulation [28, 29], which in turn simplifies their energy budgets. Accordingly, in describing the full life-cycle of loggerhead turtles, we used the least complex DEB formulation called the standard DEB model [24, 25, 26]. In this model, the state of a turtle is captured by three state variables: reserve, *E* (energy in joules, J), structure, *L* (length in centimeters, cm), and maturity, *E_H_* (J). Reserve is a maintenance-free energy buffer between the environment and the turtle that quantifies metabolic memory. Energy in reserve is readily mobilized to power metabolic processes. Structure, by contrast, is built and maintained using energy mobilized from reserve. Finally, maturity is a maintenance requiring quantity that does not contribute to body mass. It is quantified as energy that was cumulatively invested in maturation (preparation for the adult stage). Maturity controls metabolic switching (e.g., the onset of first feeding or the onset of reproduction) and, analogous to structure, is maintained with energy mobilized from reserve.

If sufficient food is available in the environment, all three state variables are increasing functions of age, yet maturity is assumed to remain constant upon reaching the adult stage. In this stage, energy previously used for maturation is redirected to reproduction. Some organisms reproduce intermittently, implying that energy is stored in a reproduction buffer. The state of the reproduction buffer is tracked using an auxiliary variable denoted *E_R_*.

Dynamics of the state variables are determined by energy flows denoted universally *ṗ*^*^ (unit J d^-1^; Figure 1):

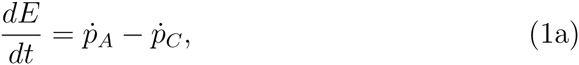

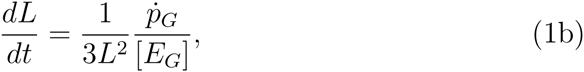

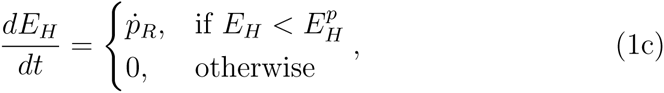

where [*E_G_*] (unit J cm^-3^) is the volume-specific cost of structure, and 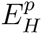 is maturity at puberty marking the beginning of the adult stage. In this stage, we replace Eq. (1c) with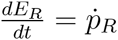. Energy flows appearing in the system of Eqs. (1) are defined as follows:

**Figure 1:**
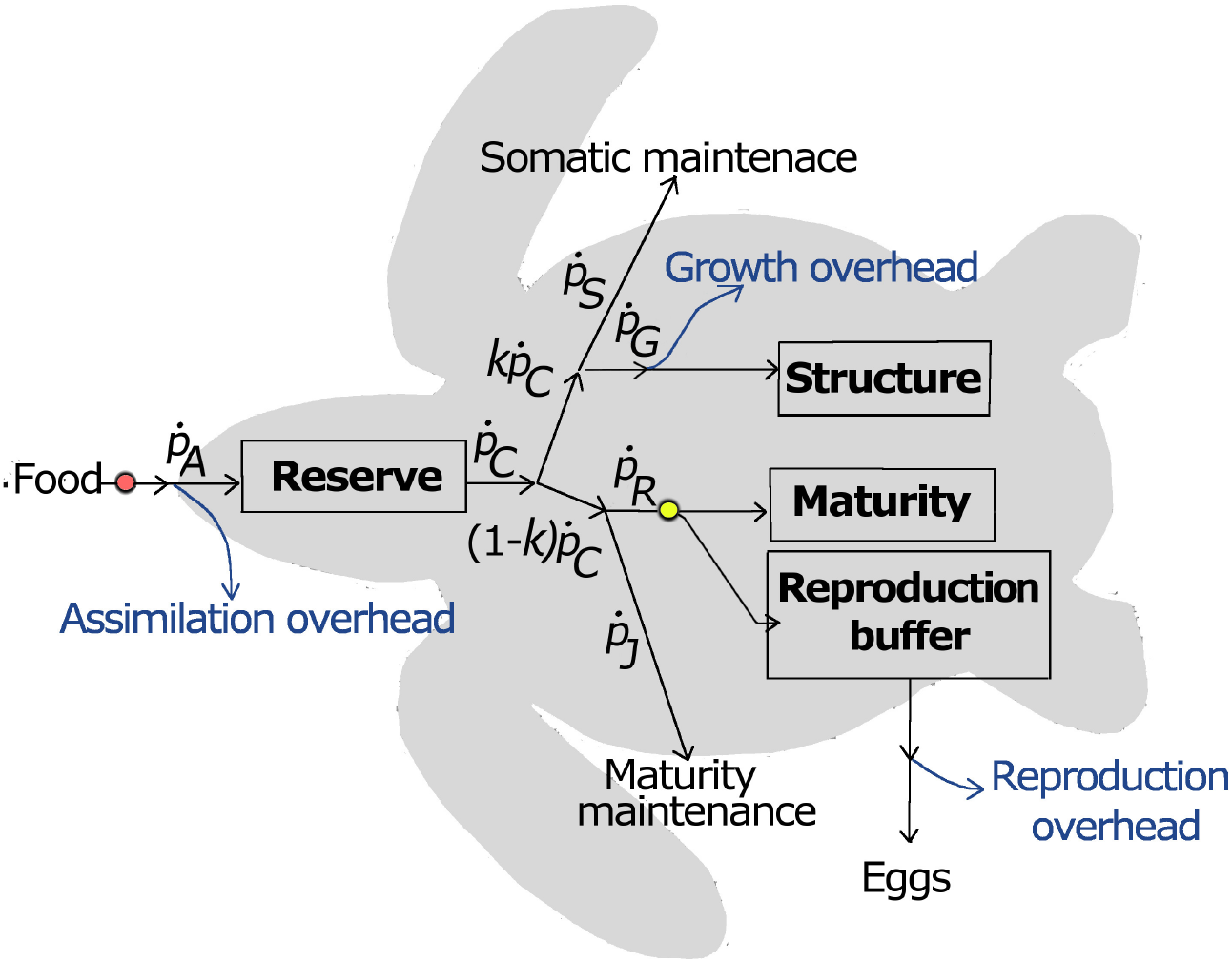
A schematic representation of the standard DEB model describing a sea turtle: Three state variables are reserve (*E*), structure (*L*), and maturity (*E_H_*). An auxiliary variable is needed to track the state of the reproductive buffer. Metabolic energy flows are: *ṗ_A_*-assimilation, *ṗ_C_*-mobilization, *ṗ_M_*-somatic maintenance, *ṗ_G_*-growth, *ṗ_R_*-maturation/reproduction, and *ṗ_G_*-maturity maintenance. The circles indicate metabolic switches that occur when a certain level of maturity is reached: the onset of feeding when 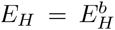 (red circle), and the onset of reproduction when 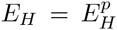 (yellow circle). Detailed definitions of these concepts are given in the main text.

**Assimilation,** 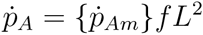, is the fraction of the daily feed ration that gets fixed into reserve, where {*ṗ_Am_*} (unit J cm^-2^ d^-1^) is the surface area-specific maximum assimilation rate and *,ƒ* is the scaled functional response equivalent to the ratio of the actual and the maximum feeding rate of an individual. The scaled functional response quantifies food availiability (i.e., *f* = 1 under unlimited food availability and *f* = 0 when food is unavailable) and in many cases can be written as

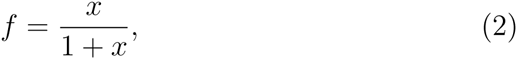

with *x* being the food density scaled by the half-saturation constant of the type-II saturating function (see p. 32 of [24] for details).

**Mobilization,** 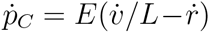, is the flow of energy mobilized from reserve to power metabolic processes, where parameter *v* (unit d^-1^) is the energy conductance and, for [*E*] = E/L^3^, the specific growth rate is

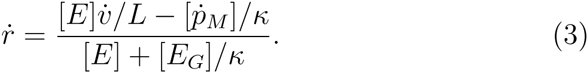

Here, [*ṗ_M_*] (unit J cm^-3^ d^-1^) is the volume-specific somatic maintenance rate. Mobilized reserve is partitioned according to the *κ*-rule: fixed fraction *κ* is allocated to satisfy the organism’s somatic needs (somatic maintenance and growth), whereas the rest is allocated to maturity maintenance and maturation (before puberty) or reproduction (after puberty).

**Somatic maintenance**, *ṗ_M_* = [*ṗ_M_*]*L*^3^ is the flow of mobilized reserve energy needed to maintain the structure of given size *L*^3^.

**Growth**, *ṗ_G_ = Kṗc — ṗ_M_*, is the flow of mobilized reserve energy invested into the increase of structure after satisfying the somatic maintenance needs.

**Maturation,** *ṗ_R_* = (1 — *ĸ*)*ṗ_c_* — *ṗj*, is the flow of mobilized reserve energy towards increasing the level of maturity (*E_H_*), after satisfying the maturity maintenance, *ṗj*.

**Maturity maintenance,** 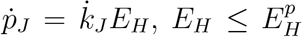, is a flow (analogous to somatic maintenance) that quantifies the mobilized reserve energy necessary to maintain the current level of maturity. Parameter k_J_ (unit d^-1^) is called the maturity maintenance rate coefficient. At the onset of the adult stage when the level of maturity reaches 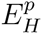, the organism starts to invest energy into reproduction instead of maturation. Hence, reproduction starts and maturity stops increasing.

All model parameters are conveniently summarized in Table 1.

Reserve and structure are abstract state variables that can be linked to commonly measured quantities such as length or body mass. A measurable length of a turtle, e.g., straight carapace length (SCL, *L*_SCL_), is related to the structural length (*L*) by the shape factor (δ_M_):

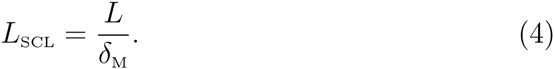

Body mass includes contributions from both reserve and structure (assumed here to have the same specific density, *d_V_* = *d_E_*). The contribution of reserve, in particular, is dependent on food availability ƒ. We have:

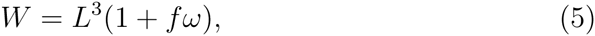

where 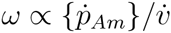 quantifies how much reserve contributes to body mass at ƒ =1. In an adult (female) loggerhead turtle, the reproduction buffer (*E_R_*) also plays a role in determining body mass [30]. However, the dynamics of this buffer were neglected because our interest lies with the overall investment of energy into reproduction rather than the detailed modeling of a reproductive season (e.g., timing and duration).

**Table 1.**
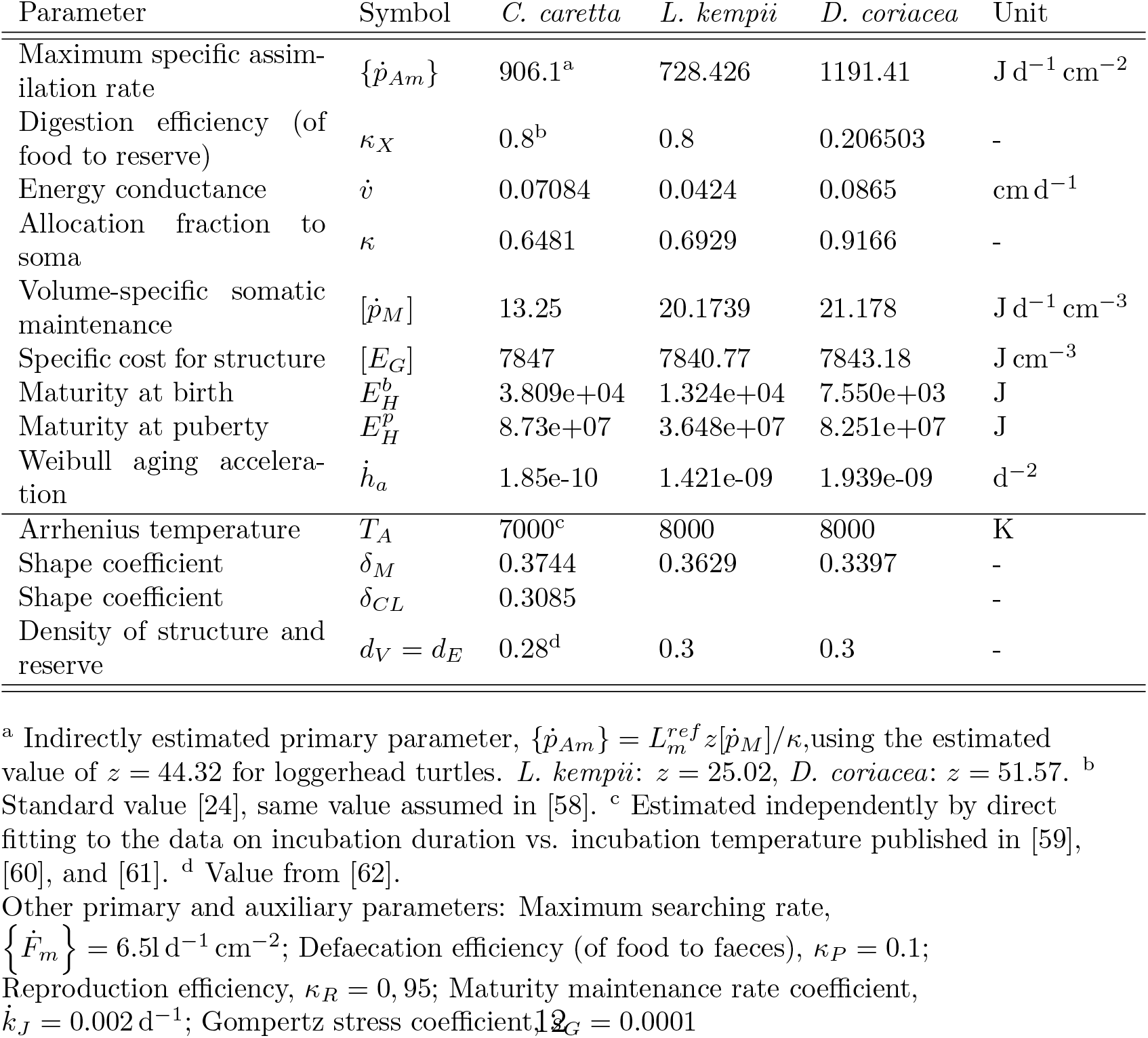
List of primary and auxiliary parameters for the North Atlantic loggerhead turtle (*Caretta caretta*) estimated using the covariation method [27] (unless specified differently). An additional shape parameter *δ_CL_* was used for the data where the type of length measurement had not been specified [54, 55]. (Preliminary) parameter values for two other sea turtles in the add_my_pet library are given for comparison: Kemp’s ridley (*Lepidochelys kempii*) [56], and leatherback turtle (*Dermochelys coriacea*) [57]. Typical values for a generalized animal with maximum length 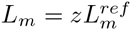. (for a dimensionless zoom factor *z* and 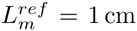), can be found in [24], Table 8.1, p. 300 and [27]. All rates are given at reference temperature *T_ref_* = 273 K, and food availability f = 0.81. Primary and auxiliary parameters for which the default values were used are listed below the table. Notation: symbols marked with square brackets, [], indicate that the parameter relates to structural volume (volume specific parameter), and symbols marked with curly brackets, { }, indicate that the parameter relates to structural surface area (surface area specific parameter). More details are available in Lika et al. [27], and the online DEB notation document www.bio.vu.nl/thb/deb/deblab/.

| Parameter | Symbol | *C. caretta* | *L. kempii* | *D. coriacea* | Unit |
| --- | --- | --- | --- | --- | --- |
| Maximum specific assimilation rate | $\{\dot{p}_{Am}\}$ | 906.1<sup>a</sup> | 728.426 | 1191.41 | $\text{J d}^{-1} \text{cm}^{-2}$ |
| Digestion efficiency (of food to reserve) | $\kappa_X$ | 0.8<sup>b</sup> | 0.8 | 0.206503 | - |
| Energy conductance | $\dot{v}$ | 0.07084 | 0.0424 | 0.0865 | $\text{cm d}^{-1}$ |
| Allocation fraction to soma | $\kappa$ | 0.6481 | 0.6929 | 0.9166 | - |
| Volume-specific somatic maintenance | $[\dot{p}_M]$ | 13.25 | 20.1739 | 21.178 | $\text{J d}^{-1} \text{cm}^{-3}$ |
| Specific cost for structure | $[E_G]$ | 7847 | 7840.77 | 7843.18 | $\text{J cm}^{-3}$ |
| Maturity at birth | $E_H^b$ | 3.809e+04 | 1.324e+04 | 7.550e+03 | J |
| Maturity at puberty | $E_H^p$ | 8.73e+07 | 3.648e+07 | 8.251e+07 | J |
| Weibull aging acceleration | $\dot{h}_a$ | 1.85e-10 | 1.421e-09 | 1.939e-09 | $\text{d}^{-2}$ |
| Arrhenius temperature | $T_A$ | 7000<sup>c</sup> | 8000 | 8000 | K |
| Shape coefficient | $\delta_M$ | 0.3744 | 0.3629 | 0.3397 | - |
| Shape coefficient | $\delta_{CL}$ | 0.3085 | | | - |
| Density of structure and reserve | $d_V = d_E$ | 0.28<sup>d</sup> | 0.3 | 0.3 | - |

<sup>a</sup> Indirectly estimated primary parameter, $\{\dot{p}_{Am}\} = L_m^{ref} z [\dot{p}_M] / \kappa$ , using the estimated value of $z = 44.32$ for loggerhead turtles. *L. kempii*: $z = 25.02$ , *D. coriacea*: $z = 51.57$ . <sup>b</sup> Standard value [24], same value assumed in [58]. <sup>c</sup> Estimated independently by direct fitting to the data on incubation duration vs. incubation temperature published in [59], [60], and [61]. <sup>d</sup> Value from [62].
Other primary and auxiliary parameters: Maximum searching rate, $\{\dot{F}_m\} = 6.51 \text{ d}^{-1} \text{cm}^{-2}$ ; Defaecation efficiency (of food to faeces), $\kappa_P = 0.1$ ; Reproduction efficiency, $\kappa_R = 0.95$ ; Maturity maintenance rate coefficient, $\dot{k}_J = 0.002 \text{ d}^{-1}$ ; Gompertz stress coefficient, $\dot{k}_G = 0.0001$

For the model to capture the whole life-cycle, we need the number of eggs produced by an adult individual. In DEB, the reproductive flow is equal to the surplus energy from flow (1 – *ĸ*)*ṗ_c_* after maturity maintenance of an adult, 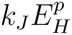, has been met:

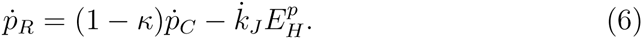

Equation (6) quantifies the investment of mother’s energy reserve into the egg production. The instantaneous reproductive output (measured in the number of eggs per unit of time) is, then, 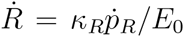, where *E_0_* is the initial energy reserve of an egg and *κ_R_* is the conversion efficiency of mother’s reserve into offspring’s reserve. Generally sea turtles produce eggs in clutches rather than continuously, and there is a trade off between clutch mass and clutch frequency [20, 30, 31]. Evolutionary constraints such as increased risks related to the nesting habitat [31, 13], mass and resource limitations, and/or metabolic heating producing excess heat that could be lethal for embryos [32, 33] all influence the clutch frequency and size. Furthermore, loggerhead turtles nesting for the first time (generally of smaller body size) produce on average half the number of clutches than those turtles that had nested previously [34]. These factors are important when energy allocated to reproduction is converted into the number of eggs per clutch (a necessity due to data availability), but do not affect the estimation of the amount of allocated energy nor the processes defining the energy budget.

### 2.2 Data used

Data on the loggerhead turtle is scarce and data sets are disjointed, meaning that studies do not share focus and methodologies can widely differ. The mechanistic nature of DEB, however, makes the assimilation of a wide variety of disjointed data types possible. Accordingly, much of the existing (published and unpublished) data could be used (Table 2). Additional information required to complete the whole life cycle has been incorporated in the model through simplifications, calculations, and/or assumptions:

- Length and body mass at puberty were calculated as the mean values of the low end of the reported size ranges for nesting females.
- The ultimate length and the ultimate body mass were calculated as the mean values of the high end of the reported size ranges for nesting females.
- Age at puberty was indirectly assumed to be equivalent to the age at first nesting and, as the age of wild nesting females is generally not known, a conservative estimate of 28 years [35, 36, 37] was used.
- Reproduction rate (R_*i*_) was assumed to be continuous (in eggs per day), rather than pulsed as in nature. This did not affect the energy balance because the total energy commitment remained the same.
- The clutch size as a function of length was calculated by assuming that:(i)the number of nests per season is the same (four) for sea turtles of all sizes (and ages); and (ii) there are no constraints on the clutch size, i.e., the clutch size is determined solely by how much energy was committed to reproduction by a nesting turtle between two reproductive seasons that are two years apart.
- The initial energy content of the egg (*E*_0_) was assumed to be the same as in green turtle eggs [38].
- The environmental (sea) temperature was assumed to be 21° C for all data relating to wild individuals, based on the average sea temperature experienced by loggerhead turtles [39]. The adults experience a higher temperature (23° C) during the nesting season [34]
- Food level was assumed to be constant, with the value approximated from the average observed ultimate size (see Table 2) and the largest observed nesting female (130cm SCL, [16]), assuming that the ratio of the two lengths corresponds to the scaled functional response, *f*, in Eq. (2).

**Table 2.** Table 2: Comparison between observations and model predictions, at the temperature that had been used for the corresponding zero-variate data (see the Section 2.2 for details), and the assumed scaled functional response f = 0.81. Values used as zero-variate data are listed in the fifth column of the table, with the corresponding relative error (’Rel. err.’) of the predictions provided in the seventh column.

| Data | Predicted | Observed range | Value used | Unit | Rel. err. (%) | Reference |
| --- | --- | --- | --- | --- | --- | --- |
| age at birth <sup>a</sup> | 52.51 | 47-60 | 57.40 | d | 8.53 | [59, 46] |
| age at puberty | 14.17 | 19-30 | 28.00 | yr | 49.39 | [35, 36, 37] |
| life span | 66.69 | >65 | 67.00 | yr | 0.46 | [40, 63] |
| SCL at birth | 5.56 | 3.9-5.06 | 4.50 | cm | 23.57 | [36, 55, 64] |
| SCL at puberty | 76.75 | 76.8-84 | 80.00 | cm | 4.06 | [45, 44, 16, 65, 19] |
| ultimate SCL | 96.35 | 98-110 | 105.26 | cm | 8.46 | [45, 44, 16, 65, 19] |
| wet mass at birth | 23.62 | 14-24 | 19.41 | g | 21.71 | [66, 59] |
| wet mass at puberty | 62.08 | 75-89.7 | 79.00 | kg | 21.42 | [16, 65] |
| ultimate wet mass | 122.82 | 148.9-180.7 | 162.62 | kg | 24.47 | [44, 65] |
| initial energy content of the egg | 209.64 | 165-260 | 210.00 | kJ | 00.17 | [38] |
| maximum reproduction rate <sup>b</sup> | 0.8556 | 0.3452-0.8630 | 0.7671 | egg/d | 11.53 | [47, 34] |

<sup>a</sup> Birth in DEB theory denotes the moment when an individual stops relying on embryonic energy reserves and starts feeding, so age at birth was calculated by summing the average incubation duration ([59, 46]), days between exiting the egg shell and exiting the nest ([46]), and days between exiting the nest and the onset of feeding (Stokes, pers.comm).
<sup>b</sup> Maximum reproduction rate was expressed as eggs per day using the number of eggs per clutch (assumed to be 140 on average), the number of clutches per nesting season, and the number of nesting seasons per year (an inverse of the remigration interval). Note that 4 clutches every 2 years, and 5 clutches every 2.5 years yield the same value of the maximum reproduction rate. The maximum reproduction rate was then calculated as $R_i = 4 \times 140 / (2.5 \times 365) = 0.7671$ .

## 3. Results

### 3.1 Model parameters and the goodness of fit

The estimated parameter values, listed in Table 1, provide a good fit between the data and the model outputs (Table 2; Figs. 2-5). In particular, life history traits such as age and length at birth, and length at maturity, are nicely reproduced by the model (Table 2). Growth curves and the relationship between body mass and length (Figures 3 and 4), as well as the relationship of clutch size to length (Figure 5) and the duration of incubation as a function of temperature (Figure 2) all agree with the data as discussed in more detail below.

**Figure 2:**
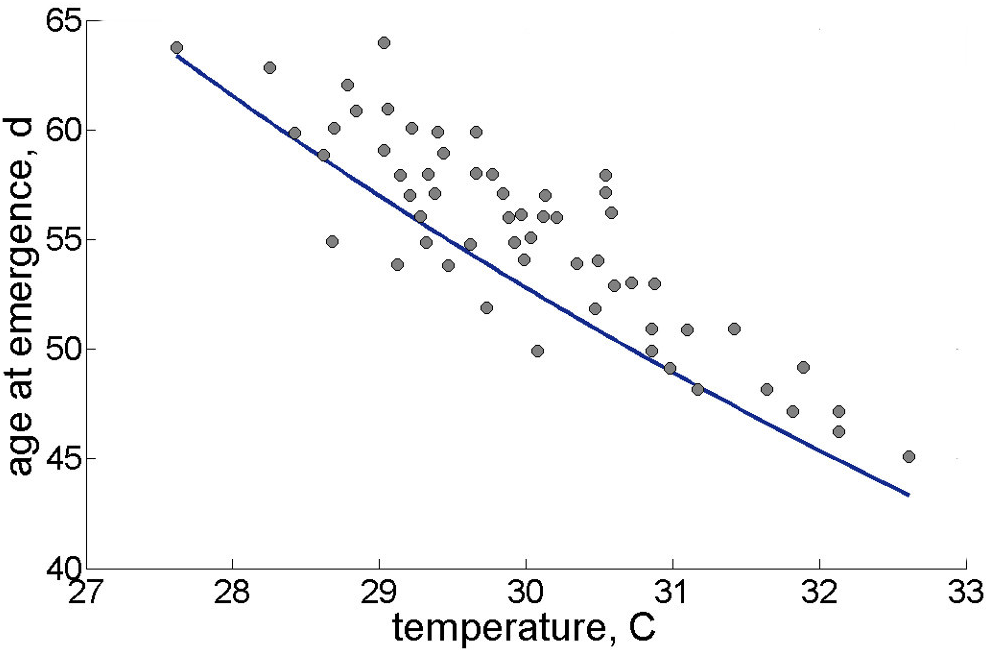
Model predictions for the duration of incubation as a function of incubation temperature. Data source: [59]; number of data points N = 61.

**Figure 3:**
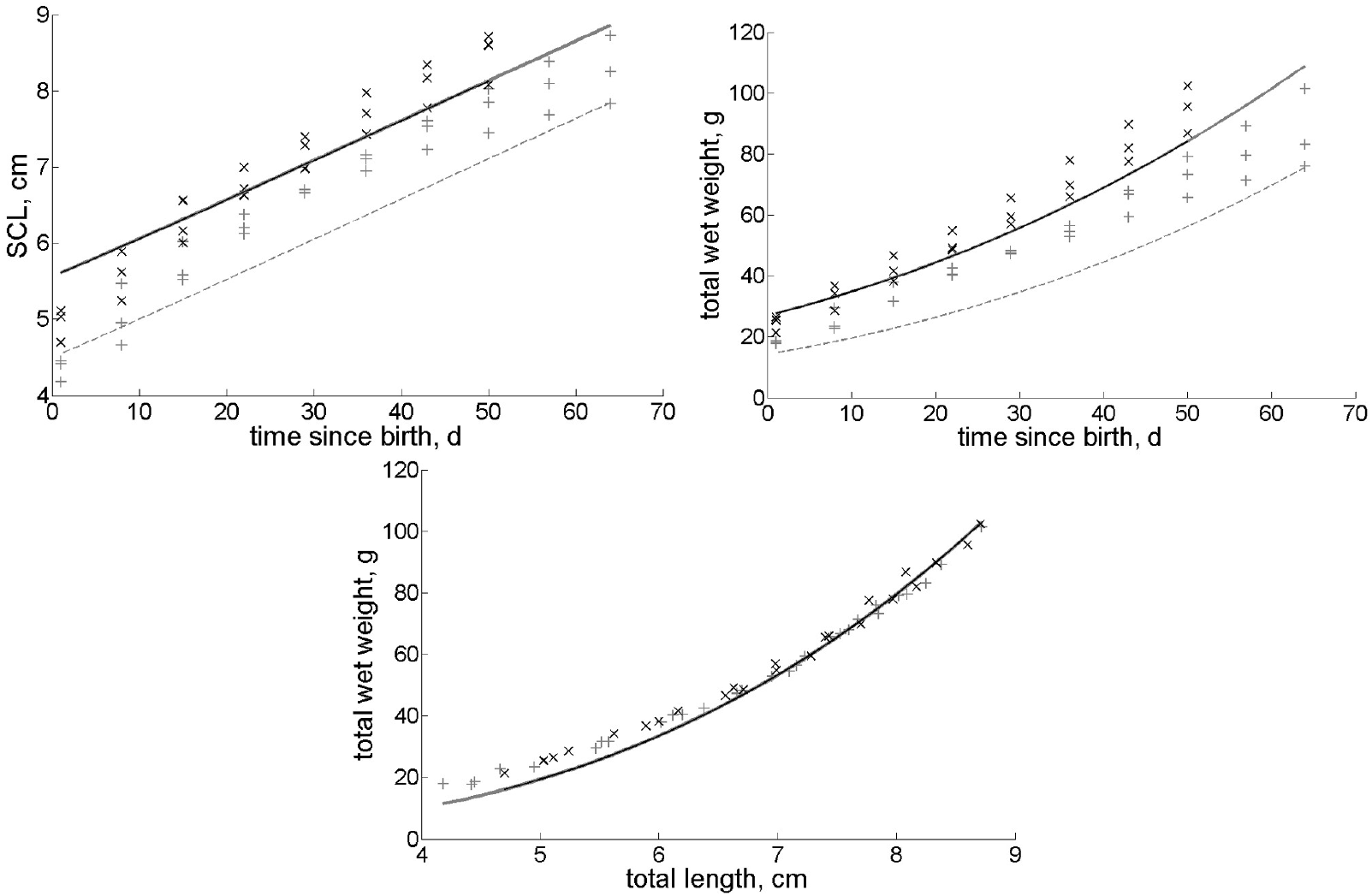
Model predictions for post-hatchlings up to 10 weeks old. Carapace length in relation to age (upper left panel), body mass as a function of time (upper right panel), and relationship of body mass and length (lower panel). Model predictions for post-hatchling growth were satisfactory when the predicted length at birth was used as a starting point (full line), but were consistently lower than the data when the observed length at birth was used to run the model (dashed line). Faster metabolism of hatchlings [67] due to their smaller size could be responsible for the underestimate. Data source: unpublished data obtained from L. Stokes. Number of datapoints: three datasets containing 10 datapoints (measurements taken weekly during 10 weeks), and three datasets containing 8 datapoints (measurements taken weekly during 8 weeks). Experimental design described in [59], and modeled as f = 0.99 and T = 27° C.

**Figure 4:**
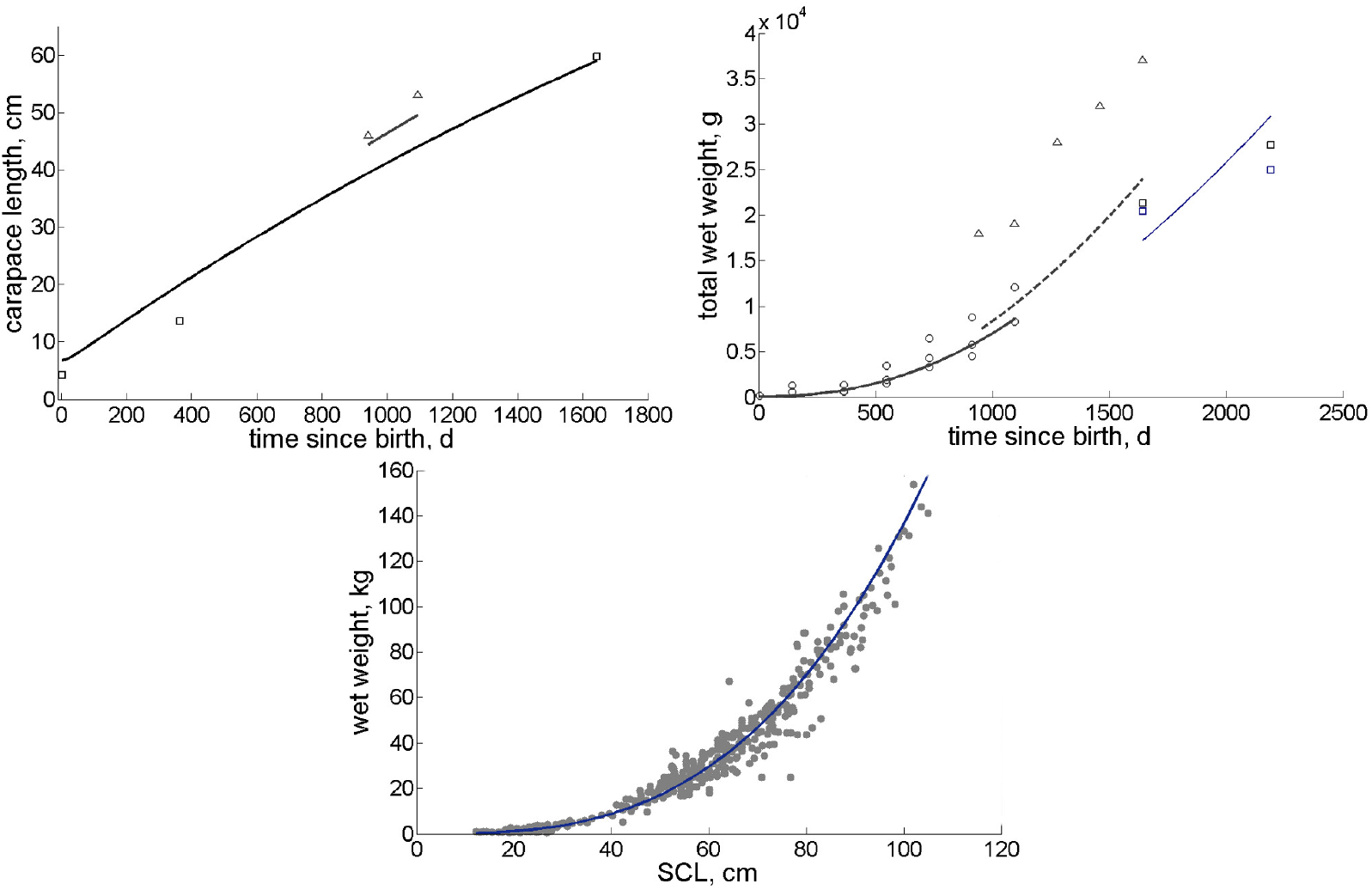
Model predictions for uni-variate data related to juveniles and adults. *Carapace length in relation to age (upper left panel)*. Data from: [54], number of datapoints N = 2 (triangles), and [55], number of datapoints N = 3 (squares). *Body mass in relation to age (upper right panel)*. Data from: [54, 64], number of datapoints N = 5 (triangles, same individual as in panel a), N = 20 (circles, three individuals); and data from [55], number of datapoints N = 4 (squares, two individuals). *Relationship of body mass and length (lower panel)*. Data from [68], number of datapoints N = 369. The exact temperature and food quantities have not been reported for some data, but most realistic results were obtained for temperature of 23° C for the fastest growing individuals (triangles in upper panels), 22° C for three individuals reared together (circles in upper right panel), and 21° C for two sea turtles reported in [55] (squares in upper left panel).

**Figure 5:**
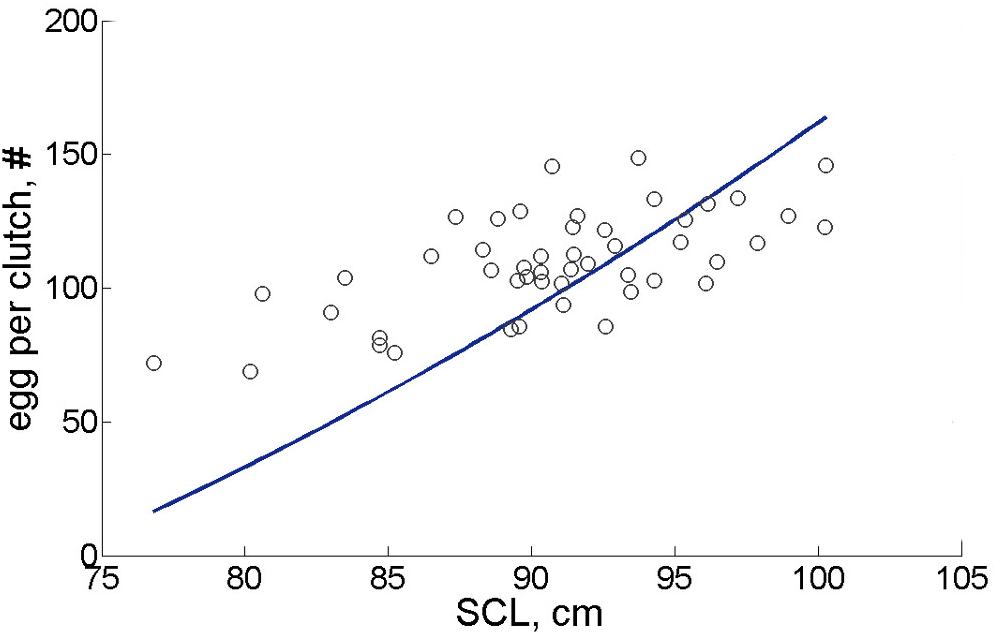
Number of eggs per clutch in relation to straight carapace length (SCL). Data from [19], number of datapoints N = 48.

Nevertheless, some traits in columns two and three of Table 2, especially the age at puberty, show apparent discord with the observations. According to the model outputs, loggerhead turtles become sexually mature at around 14 years of age, corresponding to about 76 cm SCL and 62 kg body mass. This may be because (i) the investment into reproduction precedes the first nesting and (ii) observing the exact moment at which the investment into reproduction starts is exceedingly difficult. In other words, the result is an underestimate compared to the observations deduced from size at the first reproductive event (28 years, 80cm SCL, and 79kg [17, 37, 40, 36]), yet it *is* consistent with age at puberty deduced from morphology and behavior [18, 41, 42, 43]. Other (slightly) underestimated quantities describe the ultimate size—96.4 cm SCL and 122.8 kg compared to observed 105.3 cm SCL and 162.6 kg.

Two problems arise in the context of comparisons that focus on size. First, the model estimates of body mass omit the mass of the reproduction buffer (see eq. (5)) because we assumed continuous reproduction, thus ignoring the fact that some energy (and thus mass) is stored in the reproduction buffer between two reproductive seasons. It is interesting that the cumulative (annual) wet mass of clutches produced by a turtle of 100 kg can be as much as 10 kg [30]. Accounting for this mass of the reproduction buffer would considerably decrease the current mismatch in mass between the model output and the observed values. Second, the average *ultimate* size used for parameter estimation was calculated using the high end of the reported size range from several studies. Extreme-sized individuals (that experience the best feeding conditions or that are genetically predisposed to grow large) may be introducing a bias that has a much more pronounced effect than it would have if more adults had been used for calculating the value. It is therefore encouraging that the model outputs are close to the observed average length of nesting females (92.4cm SCL, calculated from values in [16, 44, 45]) and the average body mass of adults (116.4kg [44]).

Model prediction of the incubation duration as a function of incubation temperature is quite satisfactory (Figure 2). The overall trend is correct, yet there is a small systematic bias towards the low end of the observed values. This bias suggests that although temperature explains most of the variation in the incubation duration, other factors may play an important role. Beach sand compactness and grain size, humidity, salinity of water around the nest, number of eggs in a clutch, and gas exchange of the eggs affect the incubation of loggerhead turtles as well [46, 47, 48, 49, 50], and may have to be taken into account when deducing the sex of embryos from incubation duration (e.g., [51]). In addition, metabolic heating active during the last third of the embryonic development [15, 32] could be accelerating growth and maturation (“T-acceleration”, see [52]), effectively resulting in earlier hatching and birth, and smaller than estimated size. By contrast, the previously mentioned environmental factors such as decreased respiratory gas exchange, could be prolonging the incubation [50]. The model underestimation, therefore, suggests that factors prolonging the incubation outweigh those that shorten it.

Predicted growth curves—i.e., length and body mass as the functions of age—and the resulting relationship of body mass and length are shown in Figure 3 for post-hatchlings and in Figure 4 for juveniles and adults. The carapace length estimated for *post-hatchlings* up to 65 days after birth fits the data rather well, except for a slight discrepancy for the first 10-20 days after birth. Predicted body mass during the same period fits the data even better, showing almost no discernible discrepancies. These two results suggest that the model-generated relationship between body mass and length should underestimate the data somewhat at small carapace lengths (confirmed in lower panel of Figure 3). Both the predicted carapace length and body mass of *juveniles and adults* as functions of age produce satisfactory fits over the entire period for which the data were available (Figure 4). Consequently, the relationship between body mass and length over the whole size range of juvenile and adult body sizes is in excellent agreement with the data.

Predicted reproductive output as a function of length is nearly a straight line, a result compatible with the data in Figure 5, yet the intercept and the slope of this line are respectively too low and too high. Consequently, the model predicts clutch sizes of < 50 eggs for the smallest adults and > 150 eggs for the largest adults, both of which are rarely observed in nature [47]. The predicted clutch size resulted from the conversion of energy allocated to reproduction into the clutch size—a step influenced by our assumptions on the reproductive output (see Section 2.2). However, this conversion step did not affect the prediction for the energy invested into reproduction, which is in excellent agreement with observations. The energy content of a loggerhead turtle egg is between 260 kJ and 165 kJ [38]. The predicted energy value of an egg (≈ 210 kJ) is very close to the value used for parameter estimation (2, see also [38]). Combining this value with the estimated daily energy flow to reproduction (*ṗ*_R_) of 171.34kJd^-1^ at 21° C [39], we obtain that a fully grown loggerhead turtle is capable of storing on a daily bases the amount of energy needed to build approximately one egg. If we further take the period of two years between two consecutive nesting seasons, the implication is that a fully grown (95 cm SCL) loggerhead turtle produces ≈ 595 eggs per nesting season—an equivalent of 5 clutches with 119 eggs each or 4 clutches with 148 eggs each, thus matching observations [34, 53, 38].

### 3.2 Determinants of body and energy reserve sizes

Body and energy reserve sizes are among the most important ecological parameters. Species body size, for example, positively correlates with survival [69, 70, 71] that, alongside fecundity, controls the population growth rate. The maximum structural length of loggerhead turtles, *L_m_*, is achieved for *f* = 1 and given by equation

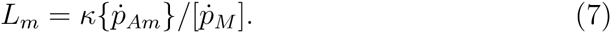

*L_m_* is determined by three parameters: allocation fraction to soma k = 0.6481, maximum surface-area specific assimilation rate {p_Am_} = 906.1 J d^-1^ cm^-2^, and the maximum volume-specific maintenance rate [*ṗ_M_*] = 13.25 J d^-1^ cm^-3^. Based on equation 7, we see that assimilation (proportional to {p_Am_}) is energy input acting to increase size (and likely survival), while maintenance (proportional to [*p_M_*]) and reproduction (proportional to (1 — κ)) are unavoidable energy outputs with the opposite effect. These parameter values in conjunction with shape factor *δ_M_* = 0.3744 correspond to the theoretical maximum carapace length of 118 cm.

Our results indicate that, on the one hand, loggerhead turtles reduce the attainable maximum size from 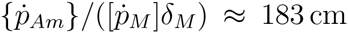 (for *k* = 1) by investing (1 – *k*) ~ 35% of the mobilization energy flow into reproduction, to already mentioned 118 cm. On the other hand, this same investment permits that an energy equivalent of approximately one whole egg at f = 0.81 and almost two eggs at f = 1 is set aside on a daily basis. The investment of energy into reproduction controls fecundity and is particularly important as one of the two chief determinants of the population growth rate. Does such an investment result in the optimal reproductive output? It turns out that at estimated k = 0.6481, the largest adults achieve only 33% of the optimum of around 6 eggs per day at f =1 (Figure 6). Achieving the optimum requires k = 0.3522. We thus find that the reproductive output of loggerhead turtles is suboptimal. A possible reason is that improved reproduction at lower k fails to offset the negatives (lower food assimilation and lower survival) associated with smaller carapace length.

**Figure 6:**
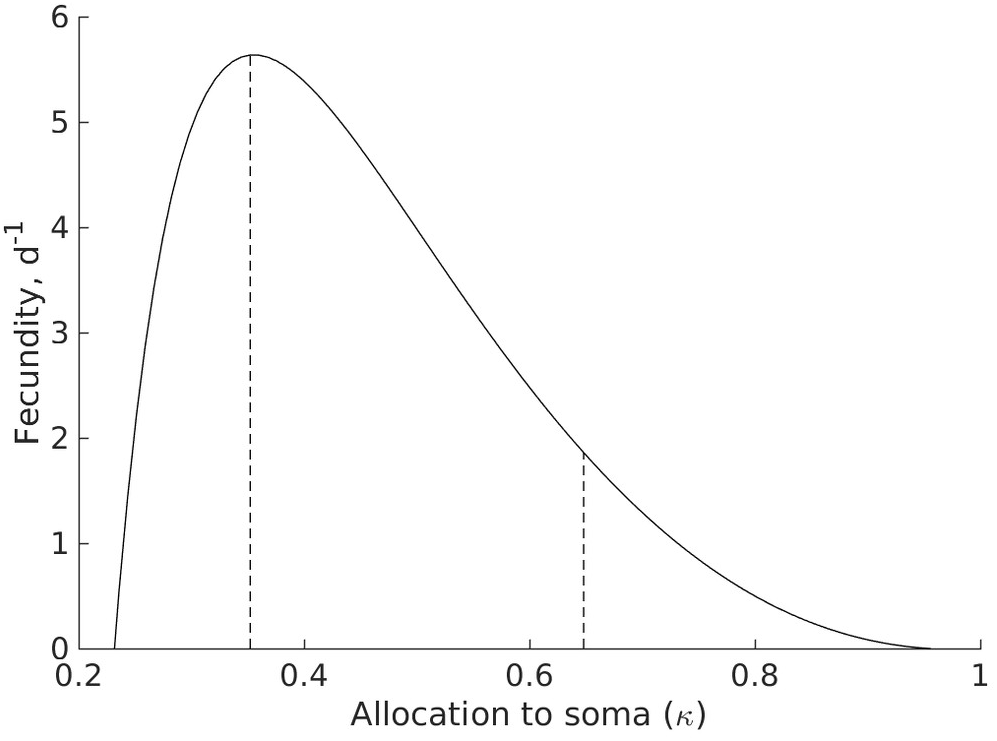
Maximum egg production of the largest loggerhead turtles (eq.2.58 in [24]) as a function of allocation to soma (parameter κ), at *f* = 1. Egg production at estimated *κ* = 0.6481 is suboptimal and amounts to only 33% percent of the optimum at κ = 0.3522. By sacrificing body size to increase the investment into reproduction (lower κ), loggerhead turtles have the potential to nearly triple their egg production. A possible reason why production remains suboptimal is that the benefit of higher fecundity (that would lead to higher population growth rate) fails to offset the negatives of smaller carapace length (that decreases the population growth rate via lower survival).

Energy in reserve is another ecologically important parameter because it indicates how well a species can endure low food availability. The ability to maintain structure in starvation is best represented by energy density, [E], the size of reserve relative to structure: [E] = *E/L_3_*. Maximum energy density, 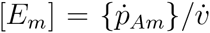, for a loggerhead turtle amounts to 12791 J cm^-3^. At maximum food availability (*f* = 1), reserve comprises 66.5% of body mass, whereas at more realistic *f* = 0.81, the percentage slightly decreases to 61.7%. In either case, the relative contribution of reserve to body mass is very large, suggesting that loggerhead turtles handle starvation rather well.

One indicator of how well an organism fares under starvation is the time to reserve depletion, *t*_†_. While there is no single general recipe for how organisms handle starvation within DEB theory (see [24], Section 4.1), the starvation mode starts when the mobilization flow, ṗ_c_ is unable to satisfy somatic maintenance according to the kappa rule, i.e., when *ĸṗ_c_* = *ṗ_M_* and hence 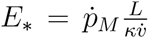. Then the special rules for starvation are applied until energy reserve is completely depleted. The time to depletion depends on the size of the individual, as well as on the strategy for handling starvation (Figure 7). While the estimates of *t*_†_ may not be completely accurate, they serve as a good qualitative measure of starvation ability. First, larger individuals have more time before experiencing problems due to unfavorable feeding conditions (Figure 7). Second, the reserve size of loggerhead turtles is such that it provides a substantial buffer against variable food availability in the environment. Even mid-sized individuals at about 50 cm carapace length have enough energy in reserve that it takes a full year before this energy is depleted. The potential to bridge long gaps in feeding might be a trait shared with other sea turtle species as indicated by the ability of sea turtles to easily sustain prolonged periods of little or no feeding during energetically demanding reproductive seasons [72].

**Figure 7:**
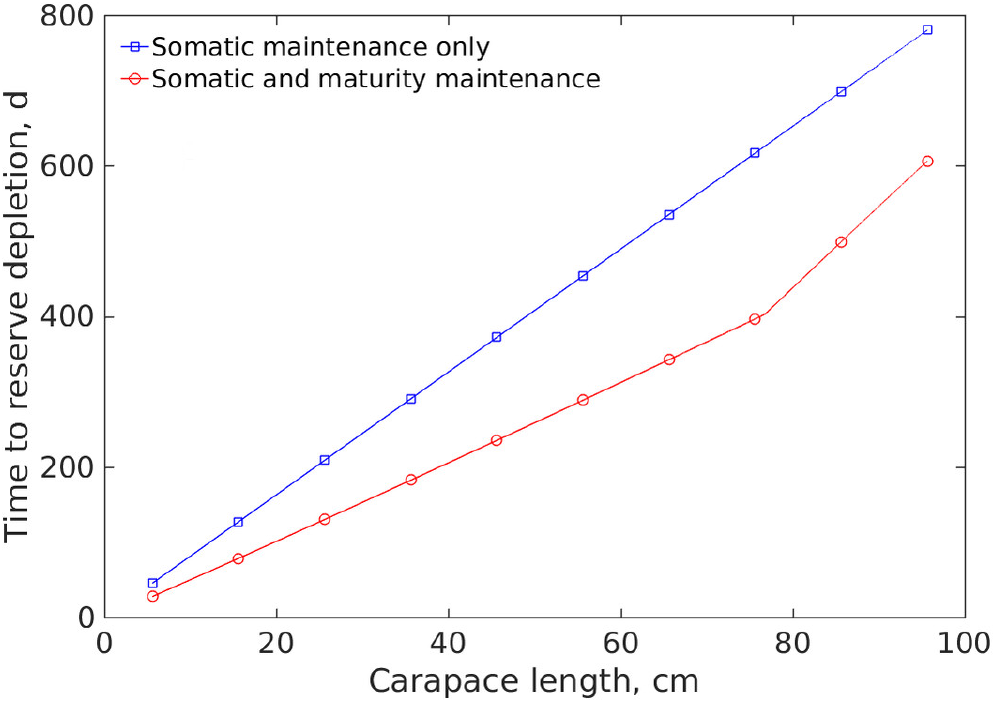
Time to reserve depletion, if, as a function of carapace length. Two possibilities are considered: (I) energy is mobilized only for somatic maintenance, 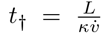 (blue squares) or (II) energy is mobilized for both somatic and maturity maintenance: 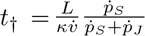 (red circles). Although larger individuals take more time to deplete their energy reserve, loggerhead turtles of any size should be able to tolerate substantial variability in feeding conditions, including prolonged periods of starvation.

## 4. Discussion

We successfully reconstructed the energy budget of loggerhead turtles using preexisting—scarce and disjointed—datasets. Such a reconstruction adds value to the data through new insights into physiology and ecology of the studied species, without additional empirical work. Gaining these new insights became possible only after jointly considering all the data within the unifying framework of DEB theory. Our unifying approach thus complements empirical studies that by necessity have a narrower focus.

Among the successfully reconstructed aspects of the energy budget, we first look at the embryonic development. The value of parameter 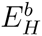 indicates that embryos on average spend 37 kJ of energy for maturation. How does this value compare with measurements? The total measured energy available at the beginning of the embryonic development (i.e., the energy of an egg) is around 210 kJ [38], whereas the total energy of hatchlings with the yolk sac at birth is around 125 kJ (calculated using measurements in [62]). The difference of 85 kJ between these two empirical values is in reasonable agreement with 62 kJ measured independently by respirometry [60] and represents the energy dissipated by embryos. A comparison between the value of 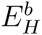 (37 kJ) and empirically determined dissipation (62-85 kJ) suggests that embryos roughly use anywhere between 40 to 60% of dissipated energy for maturation, while the rest is distributed between maintenance and growth overheads (see also Figure 8). Important in this context is the fraction of the initial reserve still left at birth because it is one of the main factors determining the resilience of hatchlings during their migration to the feeding grounds. At *f* = 0.81, for example, hatchlings have about 35 days until reserve depletion (Figure 7), assuming that the parameters remain constant throughout the ontogeny.

**Figure 8:**
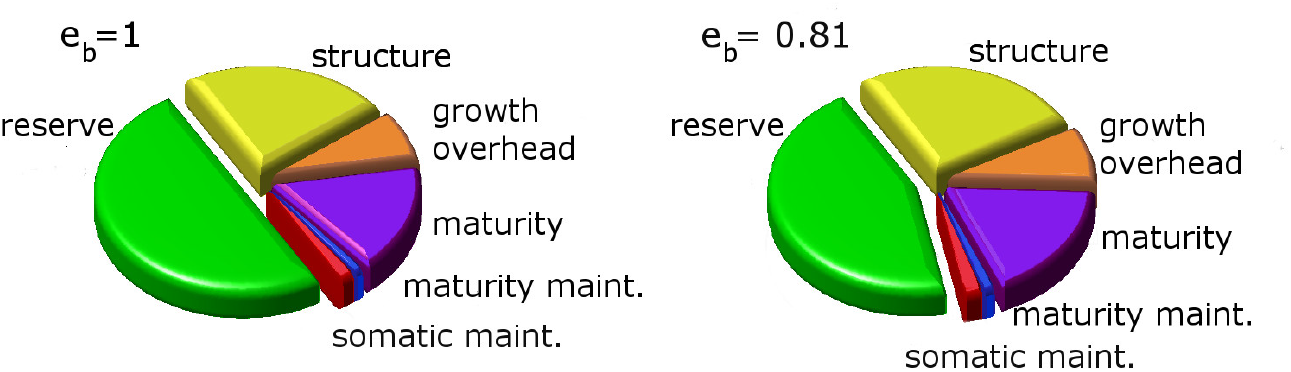
Cumulative energy investment during embryonic development, plotted at two food availabilities (*f = *e_b_* = 1 and *f* = *e_b_* = 0.81*). The lower food availability is experienced by the North Atlantic loggerhead population. If food availability were high (f = 1), about half of the initial reserve would have been dissipated into the environment or consumed for the growth of structure before birth, whereas the remaining half would still have been available to hatchlings after birth. In reality, less than half of the initial reserve is left at birth. The exact fraction is important for further development and survival because the size of the remaining reserve (partly visible as the external yolk sac) determines, e.g., the period that hatchlings survive before reaching the feeding grounds.

Among the basic DEB parameters listed in Table 1, four are expected to predictably scale with the maximum size of a species ({*ṗ_Am_*}, 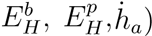, while the rest are expected to remain rather constant [24]. This scaling property can be used to further reaffirm the consistency of estimated parameter values, which we exploit by making comparisons with related species. Preliminary estimates of the standard DEB parameters were available in the online add_my_pet library [73] for two other species of sea turtles, Kemp’s ridley (*Lepidochelys kempii*) [56] and leatherback turtle (*Dermochelys coriacea*) [57]. The value of the maximum surface-area-specific assimilation rate ({*ṗ_Am_*}) falls within the range of values defined by these two species (Table 1), which is expected because loggerhead turtles are larger than Kemp’s ridley, but smaller than leatherback turtles [35]. However, both maturities (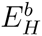 and 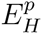) are higher and the aging acceleration (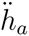) is lower for loggerhead turtles than for the other two species. While these mismatches make us cautious, they are also encouraging in the sense that the orders of magnitudes of the parameter values are similar, suggesting that the preliminary estimates for Kemp’s ridley and leatherback turtle can be greatly improved with the inclusion of more data.

The surface-area-specific maximum assimilation rate, {*ṗ_Am_*}, is determining how much energy will be assimilated into the energy reserve. The size-dependent energy budget relative to energy assimilation visualized in Figure 9 provides insight into the changes in allocation throughout the ontogeny of the loggerhead turtle (at ƒ = 0.81), and can be used as a powerful tool for exploring additional implications of changes in food availability. The proportion of assimilated energy remaining in energy reserve, as well as the energy allocated to growth, gradually reduce with size (Figure 9) as a direct consequence of the fact that most energy flows (e.g., mobilization, somatic and maturity maintenance) scale with structural volume, *L*^3^, while the assimilation scales with structural surface area, *L*^2^. Furthermore, in an energy budget of a fully grown individual the processes of (somatic and maturity) maintenance add up to become over 3/4 of the daily budget, at which point the difference between the energy assimilated into energy reserves and that mobilized for other metabolic processes reduces to practically zero. Keeping in mind that only after the cost of maintenance has been paid can juveniles grow and fully grown adults can allocate to reproduction, our results suggest that a lower amount of assimilated energy (as a result of, e.g., lower food availability), could have drastic consequences on the growth of juveniles, and the reproduction of fully grown adults. Reproducing while experiencing lower food availability could also have consequences on the survival of post-hatchlings, as the amount of energy reserves left after embryonic development is dependent on the food availability experienced by the mother (Figure 8), and will determine how long a turtle can survive before it needs to start feeding (Figure 7).

**Figure 9:**
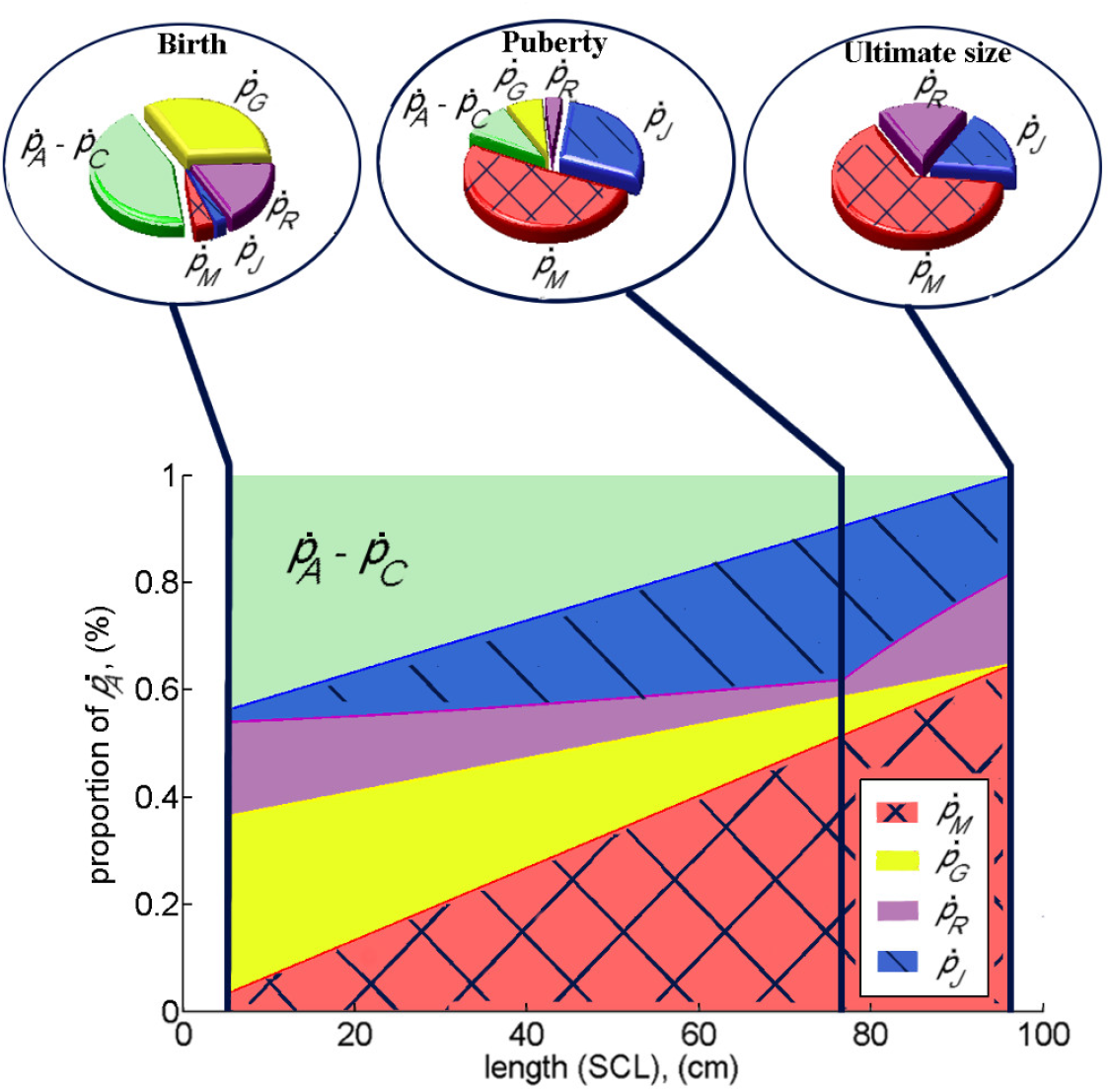
Visualization of the energy budget as a function of size. Shown are the contributions of all metabolic processes (i.e., energy flows) relative to assimilation. Special attention is given to three energetically important moments: birth, puberty, and ultimate size. Flows are calculated using the estimated parameter values for North Atlantic population (Table 1) with the scaled food availability of *f* = 0.81 experienced in the wild.

Having precise energy ingestion rates through feeding would ultimately allow various model applications such as (i) assessing the energy requirements of loggerhead turtle individuals reared in captivity [8] or (ii) investigating the ecological interactions between loggerhead turtle populations and their prey. To study the ingestion rates, we need to look into the surface-area-specific maximum ingestion rate, {*ṗ_Xm_*}, determined by the relationship

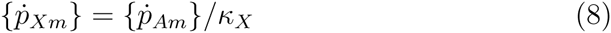

where *k*_x_ is a constant called assimilation efficiency. However, establishing the reliability of estimates of {*ṗ_Am_*} and k_x_ is difficult. Looking into the first parameter, {*ṗ_Am_*}, in more detail, we see that it determines the ultimate size of an individual (see Eq. (7)). Assuming a constant allocation to soma (*κ*) the same maximum size can be predicted with different values of {*ṗ_Am_*} and [*ṗ_M_*] as long as their ratio is constant. Our estimate of the volume-specific somatic maintenance rate for the loggerhead turtle of [*ṗ_M_*] = 13.25 J d^-1^ cm^-3^ (considerably lower than the estimates of around 20 J d^-1^ cm^-3^ for the other two sea turtle species) should be used with caution: if the estimate of [*ṗ_M_*] is too low, we may also end up underestimating the surface-area-specific maximum assimilation rate, {*ṗ_Am_*}, yet fail to recognize this underestimate as the predicted maximum size remains the same. An independent and more reliable estimate of {*ṗ_Am_*} is possible only if the precise measurements of both ingestion rates and assimilation overheads are available [74] (see also Section 11.2 of [24]). Independently estimating the value of *κ_X_*—the other parameter determining the ingestion rate—is particularly difficult because quantifying ingestion and assimilation overheads requires knowing (i) egestion, (ii) excretion, and (iii) specific dynamic action [24, 74]. Such a comprehensive set of measurements on loggerhead turtles is unknown to us, leading to the conclusion that reliable estimates of ĸ_X_ or {*ṗ_Am_*} are not possible at this moment. Hence, our estimates of the ingestion rate should be used with caution.

The only attempt to estimate a (static) energy budget of loggerhead turtles in absolute terms known to us is by Hatase and Tsukamoto [58]. The authors considered that oceanic adults of 70 kg body mass feed on energy-sparse plankton of genus *Pyrosoma,* while neritic adults of 90 kg body mass feed on energy-dense clams. Due to difficulties in obtaining precise measurements, the authors were forced to make a number of *ad hoc* assumptions to arrive at a daily energy intake of 28 454kJ (14.4 kg) of neritic food. This intake, however, seems to be too high. First, observations suggest that the feeding rate of loggerhead turtles is probably much lower: measurements of food intake by loggerhead turtles, ranging in size between 2 and 60 kg and fed anchovies in captivity, yielded a regression equation that at 20 °C gives 3.3 kg of food ingested daily when extrapolated to the size of neritic adults [75]-only about 23% of the estimate by Hatase and Tsukamoto [58]. Second, daily energy intake is unlikely to be higher than that of a species known for high energy consumption and even higher food intake. A validated energy budget exists for such a species: Pacific bluefin tuna *(Thunnus orientalis*) [7, 8, 76]. If we compare the daily energy intake of an individual Pacific bluefin tuna with the same structural size as neritic loggerhead turtle adults, it turns out that the tuna consumes about 3 400 kJ or approximately 8 times less than the value from Hatase and Tsukamoto [58]. Third, the huge intake assigned to loggerhead turtles, with a large proportion needed to satisfy the assumed basic metabolic needs, seems even less likely when put in perspective with measured or estimated metabolic rates. The neritic-sized loggerhead tur-tles routinely dissipate up to 97% less energy (extrapolated from values in Ref. [67]) than the Pacific bluefin tuna, again with the same structural size as neritic loggerhead adults: 0.03 Wkg^-1^ compared to 1.18 Wkg^-1^ at 20°C. This makes the 800% higher energy need estimated for neritic loggerhead turtles by Hatase and Tsukamoto [58] highly unlikely. It is interesting to mention that our model predicts dissipation of 0.11 Wkg^-1^ for neritic adults at 20°C with an assumed *κ_X_* = 0.8. This value drops to 0.08 Wkg^-1^ in fasting individuals, which is in line with measurements of 0.05 Wkg^-1^ by Lutz et al. [77] performed on smaller resting loggerhead turtles at 20 °C.

Estimates of energy investment into reproduction (*ṗj* and *ṗ_R_* in DEB, see Figure 1) also show a mismatch when comparing our model outputs with calculations reported by Hatase and Tsukamoto. Integrating energy invested into the reproductive branch (maturity maintenance + egg production) over two years gives an estimate of approximately 300 MJ (127MJ for maintenance, and 147 MJ for egg production) at the temperature of 23° C (the average temperature experienced by adult loggerhead turtles [39, 58]). This is markedly smaller than 1003 MJ calculated for the smaller oceanic adults nesting every second year [58], and approximately 30% less than the reproduction costs calculated for neritic Pacific loggerhead turtles nesting *every* year (435MJ, [58]). We did not separately model the neritic and oceanic adults, nor explicitly include the different expenses of migration that these two groups of adults have. However, the realistic number of eggs predicted by our model (see section 3.1) suggest that our estimate of the energy investment into reproduction is realistic.

Not all aspects of the energy budget of loggerhead turtles were captured perfectly by the model, yet even deviations of model outputs from the commonly accepted knowledge are informative. For example, we estimate that in an environment with relatively constant food and temperature, loggerhead turtles start allocating to reproduction several years before reaching the currently accepted age-at-puberty based on nesting observations. The transition to adulthood might thus be happening much earlier than currently suspected, and first nesting observed might be an inadequate proxy for puberty. The definition of “puberty”, whether it is the initial allocation to reproduction or morphological changes (e.g., tail prolongation in males) or the first nesting, therefore has to be agreed upon prior to making comparisons across studies.

Furthermore, the underestimated growth of posthatchlings during the first 15-30 days after birth (Figure 3) suggests that the description in terms of fixed parameter values throughout the whole life cycle may be somewhat inadequate. One way to speed up growth in DEB theory is exemplified by the “waste to hurry” strategy [78], whereby the increase in the values of parameters directly related to the acquisition of energy ({*ṗ_Am_*}) and metabolism (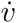 and [*ṗ_M_*]) results in faster growth, but smaller ultimate size due to a higher energetic cost. The strategy in which some energy is wasted to achieve faster growth and reduce time spent in early stages which are particularly vulnerable to predation [79] may be beneficial to post-hatchlings.

## 5. Conclusion

The standard DEB model aided the characterization of the whole life cycle of the loggerhead turtle using relatively few types of disjointed data on life-history traits and growth curves, some of which date from 1926. The estimated DEB parameter values now characterize the energy utilization patterns in the loggerhead turtle, enabling the standard DEB model to predict growth, maturation, and reproduction as a function of temperature and food (or energy reserve provided by the mother, in case of an embryo).

In addition, the parameter values enabled quantitative predictions of many energy budget features that were not (or could not be) measured directly. Examples are the plotted energy budgets at birth, puberty, and when fully grown (Figures 8 and 9). The model made it possible to study ontogeny and physiological traits such as coping with prolonged periods of starvation and the trade-offs between growth and reproduction.

Additional details could be included into the model to increase its predictive capabilities and accuracy, but whether additional predictions and accuracy warrant the increased complexity of the model highly depends on particular questions of interest. For example, precision in modeling embryonic development could be augmented by including effects of the sand (compactness, humidity, and grain size) on incubation duration and time needed from hatching to emergence. Also, metabolic heating could be incorporated into the model by increasing the temperature in simulations. Including constraints on the size and frequency of clutches, as well as explicit modeling of the reproduction buffer (as opposed to continuous reproduction), offers an opportunity to improve the conversion from allocation to reproduction (joules per day) to the reproductive output (eggs or clutches per nesting season).

The realism and precision of the model predictions could be further improved by (i) loosening the assumption that the parameters are constant throughout ontogeny, and (ii) simulating a more variable environment, reproducing some of the food and temperature variability experienced by the loggerhead turtles in the wild [12]. By allowing the parameters to vary throughout ontogeny, physiology of small loggerhead post-hatchlings can change such that temporarily increased parameter values improve growth performance, thereby reducing the risk of being eaten by predators. Simulating an en-vironment in which food availability and/or temperature drastically change might be a good approximation of the ontogenetic habitat shift when juvenile loggerhead turtles change their oceanic (colder and food poorer) environment for a neritic (warmer and food richer) one [13]. Consequently, growth curve might differ (see e.g. [5, 80, 81]) from the most commonly assumed monotonic one. Such a different environment would result also in different predictions for age at puberty. The range of observed maturation age estimates are seemingly contradictory (15-39 years, [37, 17, 40, 41, 42]).

The lower end of the range is obtained by direct observations in captivity, or deduced from morphology and behavior, while the upper end of the range is estimated using the carapace length at reproductive events. Could such a large range be explained by the time necessary to accumulate energy for reproduction after the actual maturation, or by environmental variability experienced by some loggerhead turtles in the wild?

Even without the mentioned additions and alterations, the model provides insight into physiology and ecology of the loggerhead turtle, and makes a powerful tool for conservation biology and management of sea turtles. Obtaining a set of DEB parameters for a different loggerhead turtle population (e.g., the Mediterranean population) might provide further insight into the observed [4, 19] differences in growth, maturation, and reproduction between these two populations. Information on relevant processes and life history traits (duration of life cycle phases, reproduction output, etc.) can be further studied for a range of temperature and/or food abundances to gain additional insight into physiology and ecology of the loggerhead turtle. The model is one of a full life cycle, and can be used to study the environmental effects on the physiological processes such as growth, maintenance, maturation, and reproduction. It enables exploring future scenarios, e.g., those resulting from the global climate change. In particular, the information can be used to create population models that include environmental information into the population dynamics, as it is possible to investigate how changes in temperature and food availability might affect individual physiological processes (thus affecting survival and fecundity). This is the first step toward determining the effects of environmental changes on growth and viability of a population, and the chances of success of conservation efforts.

## Acknowledgements

The authors would like to thank L. Stokes for generously sharing her data. N.M and T.K. have been in part supported by Croatian Science Foundation under the project 2202-ACCTA. M.J. was supported by the Japan Science and Technology Agency (JST) Program to Disseminate Tenure Tracking System.

